# Myeloid cell activation during Zika virus encephalitis predicts recovery of functional cortical connectivity

**DOI:** 10.1101/2023.07.06.547991

**Authors:** Shannon C. Agner, Lindsey M. Brier, Jeremy Hill, Ethan Liu, Annie Bice, Rachel M. Rahn, Joseph P. Culver, Robyn S. Klein

## Abstract

Neurologic complications of Zika virus (ZIKV) infection across the lifespan have been described during outbreaks in Southeast Asia, South America, and Central America since 2016. In the adult CNS ZIKV tropism for neurons is tightly linked to its effects, with neuronal loss within the hippocampus during acute infection and protracted synapse loss during recovery, which is associated with cognitive deficits. The effects of ZIKV on cortical networks have not been evaluated. Although animal behavior assays have been used previously to model cognitive impairment, in vivo brain imaging can provide orthogonal information regarding the health of brain networks in real time, providing a tool to translate findings in animal models to humans. In this study, we use widefield optical imaging to measure cortical functional connectivity (FC) in mice during acute infection with, and recovery from, intracranial infection with a mouse-adapted strain of ZIKV. Acute ZIKV infection leads to high levels of myeloid cell activation, with loss of neurons and presynaptic termini in the cerebral cortex and associated loss of FC primarily within the somatosensory cortex. During recovery, neuron numbers, synapses and FC recover to levels near those of healthy mice. However, hippocampal injury and impaired spatial cognition persist. The magnitude of activated myeloid cells during acute infection predicted both recovery of synapses and the degree of FC recovery after recovery from ZIKV infection. These findings suggest that a robust inflammatory response may contribute to the health of functional brain networks after recovery from infection.

**Significance Statement:** Determining the long-term cognitive impact of infections is clinically challenging. We found that the degree of myeloid cell activation correlated with the degree of recovery of functional connectivity after recovery from ZIKV encephalitis. Using functional cortical connectivity, we demonstrate that interhemispheric cortical connectivity is decreased in individuals with acute ZIKV encephalitis. This correlates with decreased presynaptic terminals in the somatosensory cortex. During recovery from ZIKV infection, presynaptic terminals recover, which is associated with recovered interhemispheric connectivity. This suggests a role for activated myeloid cells in maintenance of cognition and further supports the contribution of synapses in the cortex to functional networks in the brain, which can be detected by widefield optical imaging. These findings also suggest neuroinflammation may play a neuroprotective role in addition to aiding in local virologic control.

## Introduction

Zika virus (ZIKV) is a mosquito-borne member of the Flaviviridae family of viruses that re-emerged in the western hemisphere in 2015, most notably for teratogenic effects resulting from infections during pregnancy. However, ZIKV outbreaks throughout the world have continued to highlight neurologic consequences of ZIKV^1, 2^, indicating that, similar to other vector-borne Flaviviruses^3^, ZIKV can impact the adult central nervous system (CNS)^4–9^. There is growing evidence that long term cognitive sequelae occur after systemic ZIKV infection, even in the absence of severe illness or encephalitis^10,11^. Further, these results do not seem to be restricted to older patients, as neurocognitive decline has been also observed in adolescents^12^ and young adults^13^. Although the extent of ZIKV effects in the hippocampus has been previously studied in murine models by our laboratory and others^14–18^, little is known about its acute effects on the cerebral cortex, including the health of brain networks, and how they might contribute to neurocognitive changes after recovery from infection.

One method to characterize brain networks is examination of functional connectivity (FC), which is accomplished via evaluation of regional correlations between intrinsic neural activity that are necessary for cerebral homeostasis and cognition^19–21^. The functional magnetic resonance imaging (fMRI) community has made great strides in mapping the connections necessary for multiple brain processes using seed-based analysis to calculate the temporal synchrony between any two brain regions at rest (i.e., “resting state FC”, rs-FC)^22, 23^. fMRI of encephalitic patients offers unique challenges to the neuroimaging community because altered mental status, which is common in encephalitis patients, results in intolerance for lengthy scans necessary for functional analysis^24^. Thus, while structural MRI is used as a diagnostic tool for encephalitis, functional neuroimaging studies have not been previously done.

In contrast with fMRI, optical imaging can be performed with fewer restrictions, reduced cost and greater availability than fMRI. In animal models, an optical approach (Widefield Optical Imaging, WOI) has elucidated cortical FC networks in the mouse^25, 26^. This method also has added advantages of imaging study subjects while awake and with head fixation, increasing data fidelity and eliminating effects of sedation and motion artifacts on functional networks. Previous WOI studies in mice have shown sensitivity to disease and have elucidated local perturbations associated with glioma growth^27^, stroke^28^, and Alzheimer’s Disease (AD).^29^ This modality, however, has not been extended to animal models of neurotropic and emerging Flavivirus infections that induce long-term cognitive sequelae, which may lead to dementia^30^.

In this study, we evaluated cortical FC in an adult murine model of ZIKV infection. Eight-week-old Thy1-GCAMP6f mice were intracranially (i.c.) infected with a mouse adapted Dakar strain (ZIKV-Dak-MA) which exhibits significant tropism for the mouse brain^31^, particularly in neurons of the hippocampus and cerebral cortex^31^. FC was compared between ZIKV-Dak-MA- and mock-infected mice during acute infection and after viral clearance and recovery. Deficits in interhemispheric FC were observed in the somatosensory cortex during acute infection and improved during recovery, despite continued memory deficits observed using behavioral testing. These changes correlated with alterations in the presynaptic marker density, synapsin, which also recovered after resolution of infection. Although myeloid cell activation did not appear to correlate with region-specific FC during acute infection, the degree of persistent myeloid cell activation during recovery did correlate with the degree of recovery of FC. Although cortical brain networks appeared to be intact after recovery, severe hippocampal injury, with associated impairment in spatial learning and memory persists after recovery from ZIKV-Dak-MA infection. These data indicate that cortical networks may recover in the setting of subcortical injury, but this does not contribute to recovery of hippocampal-based learning tasks. Importantly, persistent myeloid cell activation plays a central role in the restoration functional brain networks long after their role in virologic control is completed.

## Results

### Infection of Thy1-GCAMP6f mice with ZIKV-Dak-MA leads to destruction of hippocampus with loss of spatial learning ability

Previous data demonstrated that ZIKV targets the hippocampi of adult mice after i.c. infection with an African strain of ZIKV from Dakar, Senegal (ZIKV-Dak)^14^. Recent data demonstrated more severe CNS infection using a mouse-adapted strain of ZIKV-Dak (ZIKV-Dak-MA)^31^. Prior studies using WOI have demonstrated the ability to detect disrupted functional networks in the setting of subcortical injury and given the extent of ZIKV-Dak-MA infection in the hippocampus and throughout the cortex, we hypothesized that WOI would allow us to assess functional cortical network disruption by ZIKV. To determine the extent of infection and cognitive correlates of this more severe ZIKV brain infection, eight-week old C57Bl/6J mice were intracranially (i.c.) inoculated with either 5% fetal bovine serum in phosphate buffered saline (i.e,,mock), ZIKV-Dak, or ZIKV-Dak-MA. Survival, weight loss, and encephalitis scores of mice infected with ZIKV-Dak-MA were similar to that seen in mice infected with ZIKV-Dak (Supp. Fig 1A-C). We also found that 8-week-old C57Bl/6J mice had similar levels of ZIKV-Dak-MA viral replication in the olfactory bulb, hippocampus, cortex and cerebellum (Supp. Fig 1D-G) compared to 4- to 5- week old C57Bl/6J mice, published previously^31^. Using *in situ* hybridization to assess viral RNA, we found that ZIKV-Dak-MA exhibited tropism for the hippocampus, similar to ZIK-Dak^14^, but also infected the cerebral cortex at 7 days post-infection (dpi) (Supp. Fig 1H,I). Evaluation of spatial learning via the 5-day Barnes maze spatial learning paradigm (as described previously^32^) at 42 dpi revealed more significant numbers of errors in ZIKV-Dak-MA- versus ZIKV-Dak-recovered mice (Supp. Fig. 1J). ZIKV-Dak-recovered mice were slower to learn the maze than mock-infected animals but were eventually able to learn it. In contrast, ZIKV-Dak-MA-recovered animals were completely unable to learn the spatial learning paradigm. There were no significant differences between ZIKV-Dak- and ZIKV-Dak-MA-infected mice in open field testing, which measures anxiety by evaluating the proportion of time the mice spend in the center of the box compared to time spent in the periphery^32^ (Supp. Fig. 1K). In addition, no significant differences in mobility were found, which is tested in the open field test by quantifying the number of lines crossed in the demarcated box. These data suggest ZIKV-Dak-MA induces complete loss of spatial learning ability after recovery from acute infection.

### Neuronal injury and inflammation in the hippocampus during acute ZIKV infection persist after recovery

To understand the cellular correlates of ZIKV-Dak-MA infection and the significant spatial learning deficits noted after recovery from ZIKV-Dak-MA infection, we first assessed the hippocampus at 7 dpi. Quantitative immunohistochemical (IHC) detection of neurons (NeuN), activated myeloid cells (Iba1 positive) and activated astrocytes (GFAP positive) at 7 dpi revealed decreases in numbers of NeuN+ positive cells, and increased GFAP+ and IBA1+ cells (Supp. Figure 2A-E), indicating injury to the hippocampus and inflammation in the region of the hippocampus, even during acute infection. To determine if this injury persisted or recovered after viral clearance and recovery from infection, quantitative immunohistochemical (IHC) detection of neurons (NeuN), activated myeloid cells (Iba1 positive) and activated astrocytes (GFAP positive) at 42 dpi revealed severe injury in the cornu ammonis, region 3 (CA3) of the hippocampus with persistent inflammatory response, including increased GFAP expression, increased activated myeloid cells, and decreased numbers of neurons when compared with mock-infected animals (Supp. Fig. 2F,G). This persistent injury to the hippocampus was much more severe than previously reported for ZIKV-Dak and suggests more extensive destruction of the hippocampus results in the inability of mice infected with ZIKV-Dak-MA to learn the Barnes maze despite virologic control (Supp. Fig 1J).

### Acute ZIKV infection causes decreased cortical connectivity in the somatosensory cortex

Based on prior work using functional cortical connectivity (fc)-WOI to assess cortical regions involved in learning and memory^29^, we wondered if widefield optical imaging could detect disruption in cortical networks involved in spatial navigation and memory, including those with connections to the retrosplenial cortex, an integrative hub for spatial cognition^33^. To determine cerebral cortical connectivity during and after recovery from ZIKV encephalitis, we performed widefield optical imaging of Thy1-GCAMP6f mice that were i.c. mock- or ZIKV-Dak-MA-infected. Cranial window surgery and functional cortical imaging was performed at 7 dpi, which corresponds to peak encephalitis (Supp. Fig. 1C) and peak infection (Supp. Fig 1D-G). Calcium dynamics produced by genetically encoded calcium indicators (GECI’s) provide improved temporal resolution and more direct neural recording than downstream hemodynamics, while the latter provides an approximation for what the blood oxygen level dependent (BOLD) fMRI signal might produce in human subjects^22^. One measure of FC in the brain is homotopic connectivity, which measures FC of spatially mirrored locations in each hemisphere of the brain. Bilateral FC analysis (Fig. 1A,C left) revealed high levels of homotopic connectivity in mock-infected mice using both infraslow (0.009-0.08Hz) hemoglobin and delta (0.4- 4.0Hz) calcium frequency bands. The average bilateral FC value over the entire field of view (FOV) for both infraslow hemodynamics and delta calcium dynamics was significantly lower in ZIKV-infected mice (Figures 1B,D). Cluster size-based statistical thresholding revealed an FC deficit using delta calcium dynamics that was constrained to somatosensory regions (Fig. 1G), while there was a more global decrease in FC strength using infraslow hemodynamics that included the retrosplenial cortex (Fig. 1F). Overlay of the deficit in Fig. 1G onto an adapted Paxinos atlas-based cortical parcellation^25^ (Supp. Fig. 3) confirmed ZIKV-induced deficits in delta calcium bilateral FC clustered within somatosensory areas. Although, this study was not specifically designed or powered to evaluate sex as an independent variable, we did compare the average bilateral FC values for males and females using delta calcium and infraslow hemoglobin at 7dpi and found no significant difference for either infraslow hemodynamic or delta calcium imaging (Supplemental Figure 4). To confirm that the cortical needle insertion necessary for cranial infections would not artifactually result in FC differences, we compared bilateral FC in mice from the current study that were i.c. injected with mock solution 7 days prior to imaging and compared it to bilateral FC in mice that had received a peripheral injection 72 hours prior to imaging (N=8), Supplemental Figure 5). We found no differences in FC between mice who received i.c. compared to peripheral injection, which demonstrates that the intracranial needle insertion had no effect on FC measures.

**Figure 1:**
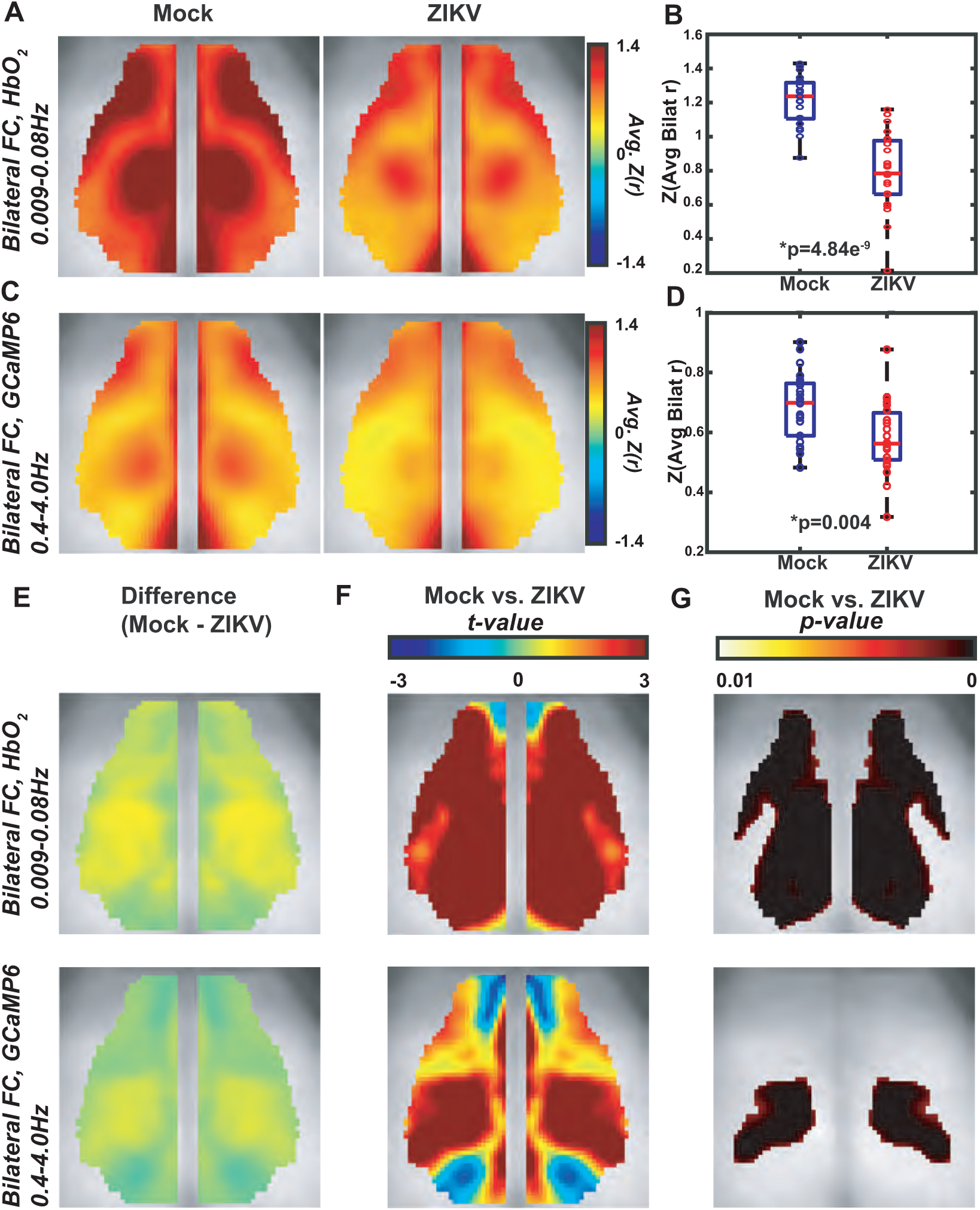
ZIKV infection reduces delta calcium and infraslow hemoglobin homotopic connectivity in the somatosensory cortex. Average (mock, N=22, ZIKV, N=23) pixel-wise bilateral FC maps across mice for A) infraslow hemoglobin and C) delta calcium. Pixel-wise difference images E) and two-sample *t*-test (left) and thresholded image (right) for *p*<0.01 by a cluster-based thresholding method for F) delta calcium and G) infraslow hemoglobin. Spatially averaged Pearson correlation value over the whole FOV for B) delta calcium and D) infraslow hemoglobin. Red horizontal bars represent the median value, while the box edges represent the interquartile range. Extending lines represent the maximum and minimum values. Significance determined by a two-sample *t*-test. Error bars are standard deviations and significance is determined by a two-sample *t*-test.

### Cortical connectivity improves after recovery from ZIKV-Dak-MA

In humans and mice, cognitive impairments may persist long after recovery from acute ZIKV infection^10, 14^. Because ZIKV-Dak-MA-recovered mice have impaired spatial learning (Supp. Fig 1J), we determined whether cortical connectivity networks measured by widefield optical imaging in recovered animals could detect alterations in cortical networks implicated in spatial learning via imaging of a cohort of mock-versus ZIKV-Dak-MA-infected mice at both 7 dpi and 42 dpi. Average (mock, N=11, ZIKV, N=8) paired bilateral FC maps using infraslow hemoglobin and delta calcium dynamics at 7dpi and 42dpi revealed complete recovery of the connectivity deficits at 42 dpi compared to 7dpi (Fig. 2). Comparison between somatosensory deficits detected at acute timepoints within a limited N=8 ZIKV dataset for delta calcium and a similar global FC deficit is present in the infraslow hemoglobin data at 7dpi via cluster extent-based thresholding (Supplemental Figures 7,8). In contrast, no deficit in either infraslow hemoglobin and delta calcium dynamics remained at 42dpi (Figure 2A,B). The bilateral maps for each mouse at 42dpi were spatially averaged across the whole FOV, as in Figure 2, and displayed as boxplots for mock and ZIKV-recovered groups (Figure 4A,B; far right columns), showing no significant differences between groups. Additionally, there were no sex-dependent significant differences in average bilateral FC at 42dpi (Supplemental Figures 4E,F).

**Figure 2.**
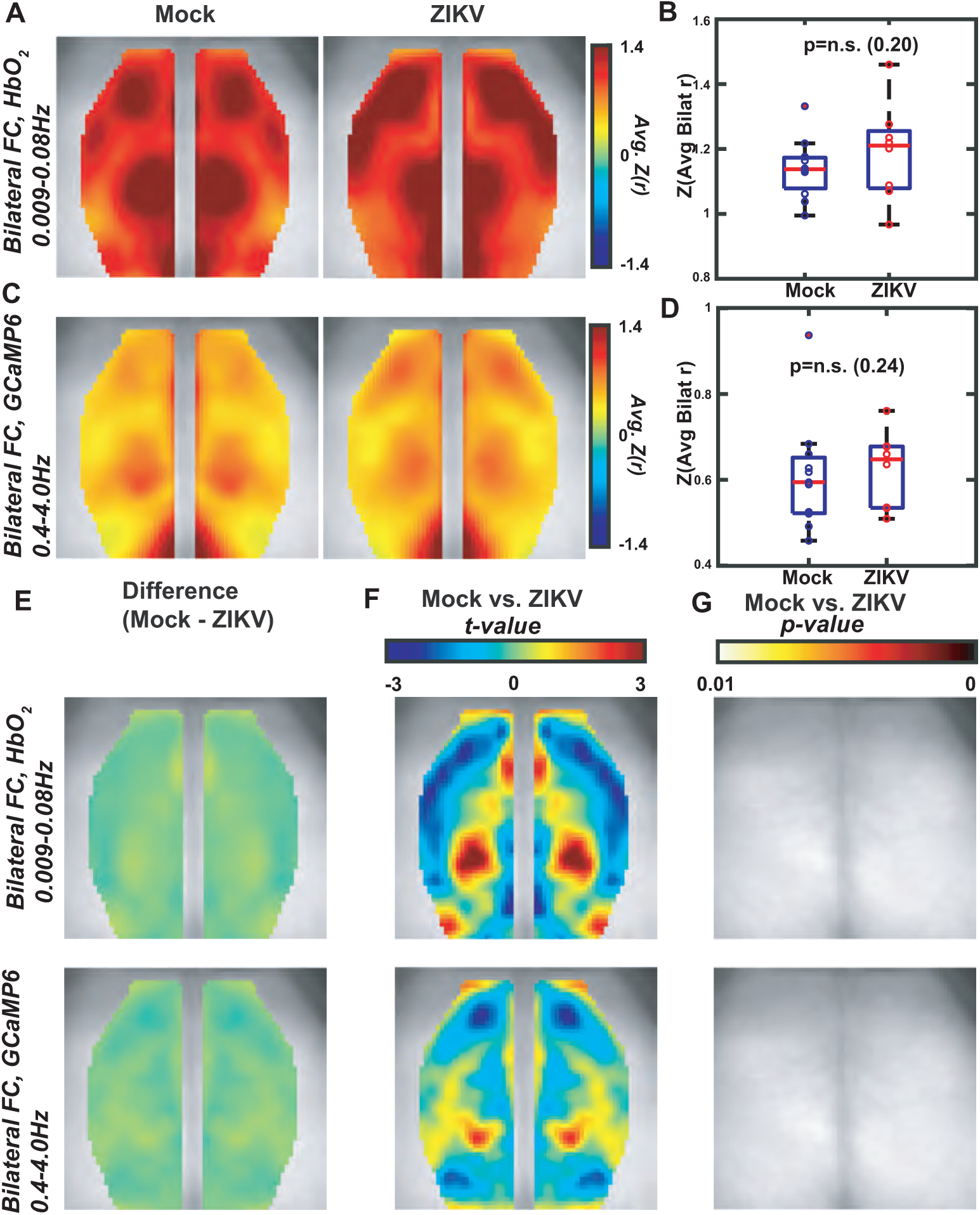
ZIKV-induced FC deficits resolve by 42dpi. Average (mock, N=11, ZIKV, N=8) bilateral FC maps at 42dpi for A) infraslow hemoglobin and C) delta calcium. Pixel-wise difference images E) and two-sample *t*-test (left) and thresholded image (right) for *p*<0.01 by a cluster-based thresholding method for F) delta calcium and G) infraslow hemoglobin. Spatially averaged Pearson correlation value over the whole FOV for B) delta calcium and D) infraslow hemoglobin. Red horizontal bars represent the median value, while the box edges represent the interquartile range. Extending lines represent the maximum and minimum values. Significance determined by a two-sample *t*-test. Error bars are standard deviations and significance is determined by a two-sample *t*-test. The one data point extending beyond the maximum value is a statistical outlier in the mock group. No significance via a paired t-test (outlier removed).

### Presynaptic termini loss during acute infection and restoration during recovery underlie alterations in FC

Prior work has suggested that human resting-state FC networks are influenced by synapses^34^. Therefore, we evaluated expression of synaptic markers that are known to be ubiquitously expressed in the cerebral cortex at 7 dpi^35^. Quantitative IHC detection of synapsin, a presynaptic marker, and PSD-95, a post-synaptic marker, within the somatosensory cortex, revealed significantly decreased detection of synapsin, but not PSD-95, in ZIKV-Dak-MA-infected mice compared with mock-infected controls (Fig 5A-D). Colocalization of synapsin and PSD95 was also decreased in mice during acute infection compared with mock-infected controls (Fig 5E), demonstrating an association between synapsin+PSD-95+ synapses and interhemispheric connectivity during acute ZIKV infection.

Because presynaptic termini loss was associated with decreases in interhemispheric connectivity at 7 dpi, we similarly evaluated expression of synapsin and PSD-95 in somatosensory cortex tissues derived from ZIKV-Dak-MA-recovered versus mock-infected animals at 42 dpi. In contrast to the decrease in presynaptic synapsin+ termini observed at 7 dpi, no differences in numbers of synapsin+ presynaptic termini or colocalization of synapsin and PSD-95 were observed between mock- and ZIKV-Dak-MA-infected animals (Fig. 5F-J). Despite recovery of synapses and cortical connectivity in mice during recovery, hippocampal injury without neuronal recovery and spatial learning deficits persist at 42 dpi (Supplemental Figure 1J). These data indicate that cortical networks, via restoration of synapses, can be restored even in the setting of severe subcortical injury.

### Alterations in cellular content are consistent with FC changes

To better understand the cellular correlates of altered FC in the somatosensory cortex at 7 dpi, we examined neurons, astrocytes, activated myeloid cells via IHC in the somatosensory cortex (Fig 3A) and compared these to a control region, the retrosplenial cortex (Fig 3B), which did not show significant changes in interhemispheric connectivity. Neuron numbers were decreased (Fig 3C,D) and GFAP+ (Fig 3E,F) and IBA1+ (Fig 3G,H) cells were increased in ZIKV-infected mice compared to mock-infected controls in both the somatosensory cortex and the retrosplenial cortex, although the statistical significance was larger in the somatosensory cortex. These findings indicate that cellular composition is associated with the FC measured by fast calcium dynamics (Fig. 3A,B). However, since infraslow hemoglobin data demonstrated that both the retrosplenial and somatosensory cortices are affected by ZIKV infection (Fig 2G), cellular content, particularly inflammation, may be driving hemodynamic signaling measured by infraslow hemoglobin imaging.

**Figure 3.**
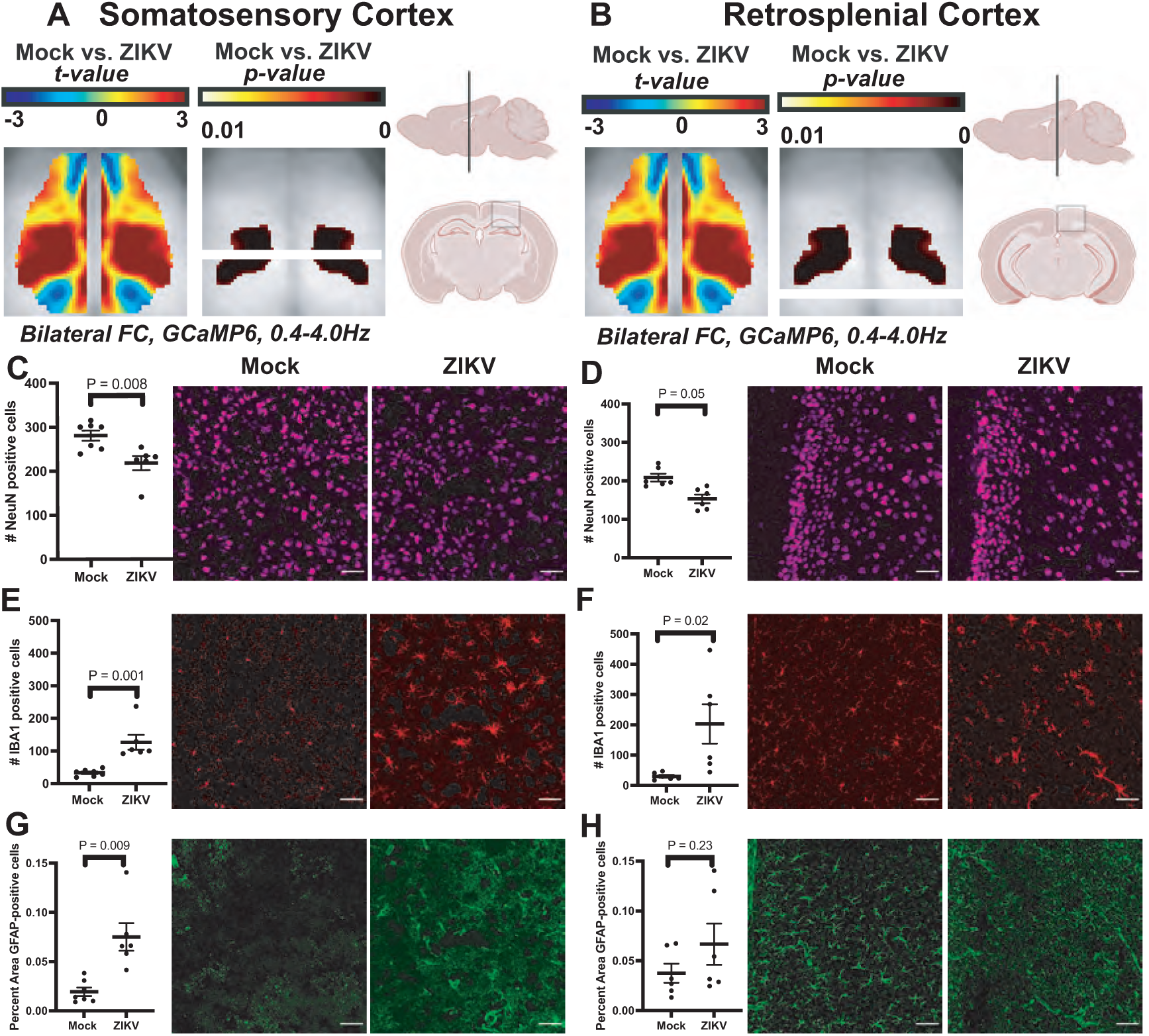
ZIKV infection induced with cellular changes not only in the somatosensory cortex, but also in a control region. (A) Diagram showing approximate location of somatosensory cortex samples in axial, sagittal and coronal planes. (B) Diagram showing approximate location of retrosplenial cortex samples in axial, sagittal and coronal planes as reference region. (C, E, G) NeuN (top), Iba1 (middle) and GFAP (bottom) staining of somatosensory cortex, where NeuN+ cells are significantly decreased, Iba1+ cells are significantly increased and GFAP+ cells are increased in Zika infected mice compared to mock mice. Mock, n=7; Zika-infected, n=6. Data were pooled from 2 independent experiments. (D,F.H) NeuN (top), Iba1 (middle) and GFAP (bottom) staining of retrosplenial cortex, where NeuN+ cells are significantly decreased, Iba1+ cells are significantly increased and GFAP+ cells trend toward increase in Zika infected mice compared to mock mice, despite no associated difference in interhemispheric connectivity. Images taken at 20x. Mock, n=6; Zika-infected, n=6. Data were pooled from 2 independent experiments. Scale bars = 50 µm.

### Myeloid cell activation supports recovery of FC and synapses after ZIKV encephalitis

To determine whether recovery of interhemispheric connectivity (Fig 3) was accompanied by reduction in inflammation, brain tissues were harvested from mice immediately following widefield optical imaging on 42 dpi, followed by IHC detection of NeuN, Iba1 and GFAP within the somatosensory cortex. While neuronal numbers in the somatosensory cortex of MA-recovered versus mock-infected mice were similar at 42 dpi (Fig 5), significantly higher levels of GFAP and Iba1 persist (Fig. 5A,B), indicating that cortical connectivity recovers (Fig 4), despite persistent inflammation in the cortex. Region-specific FC changes are calculated by comparing within-brain region parcellation of mock and ZIKV-infected mice. To determine if changes in levels of cellular markers correlate with alterations in FC, each quantity for tissues evaluated at both 7 and 42 dpi were aggregated and plotted against change in FC between 7 dpi and 42 dpi. These analyses revealed that numbers of Iba1+ cells at 42 dpi, but not NeuN+ or GFAP+ positive cells, were highly correlated with change in FC (Supp. Fig 10), indicating that persistently elevated myeloid cell activation may support recovery of functional cortical networks. When we compared FC in both mock- and ZIKV-MA-infected mice at 7 dpi and 42 dpi, we also noted that mock-infected mice had an overall trend toward decreased FC from 7 to 42 dpi, indicating a possible age-related change in FC (Supplemental Fig 11). To determine if FC at 7 dpi was predictive of FC at 42 dpi, FC values within mouse were plotted against each other, and there were no significant correlations between the two time points, indicating that 7 dpi FC was not predictive of 42 dpi FC (Supplemental Fig 10A).

**Figure 4.**
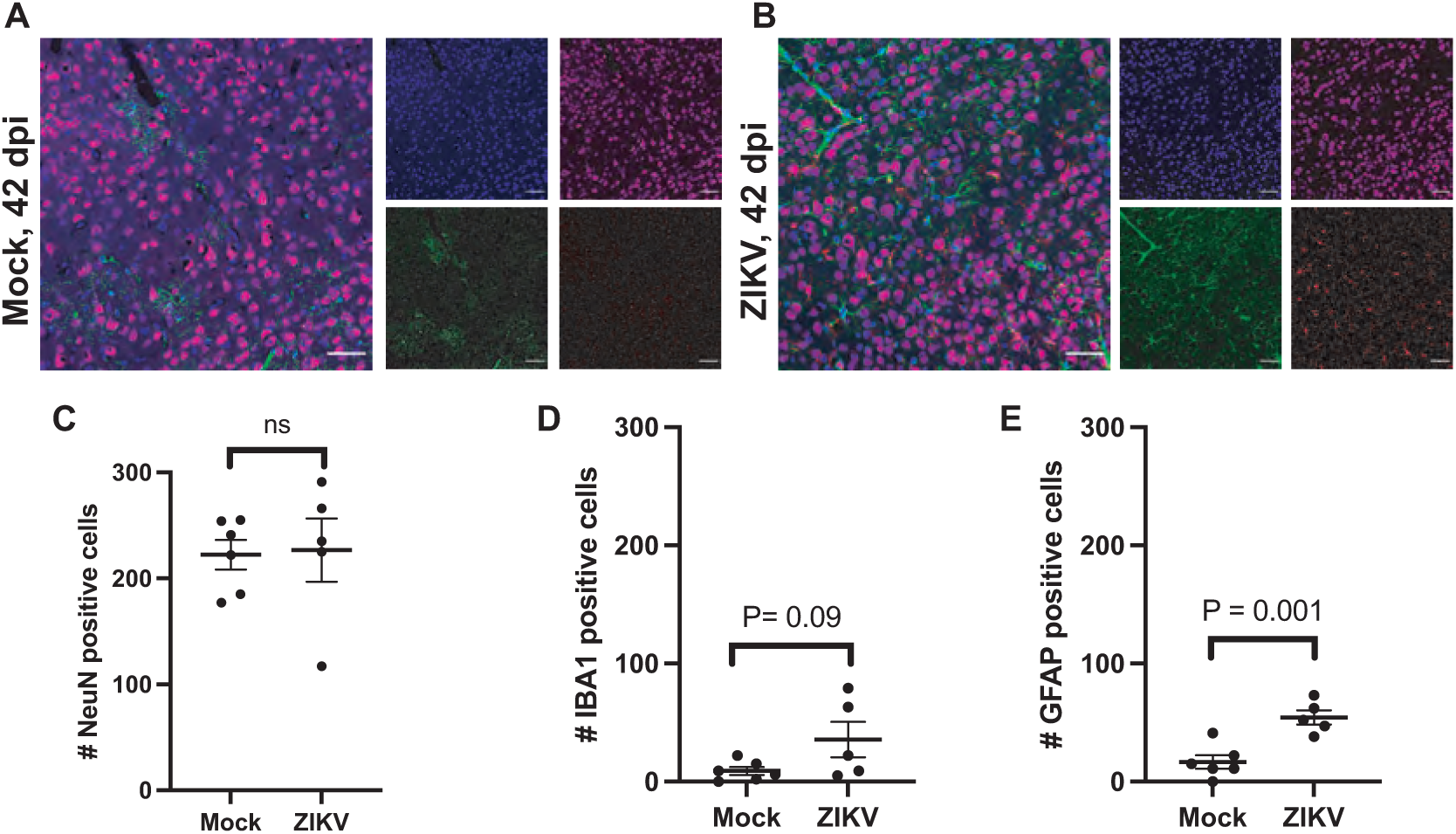
ZIKV infection resolves, but inflammation persists. Staining of mock (A) and ZIKV-infected (B) tissue at 42 dpi from the somatosensory cortex. Quantification of NeuN (C), Iba1 (D) and GFAP (E) are also displayed. NeuN positive cells appear to recover in ZIKV-infected mice, but Iba1-positive and GFAP-positive cells are increased compared to mock animals. Mock, n=6; Zika-infected, n=6. Data were pooled from 2 independent experiments. Images taken at 20x. Scale bars = 50 µm.

**Figure 5.**
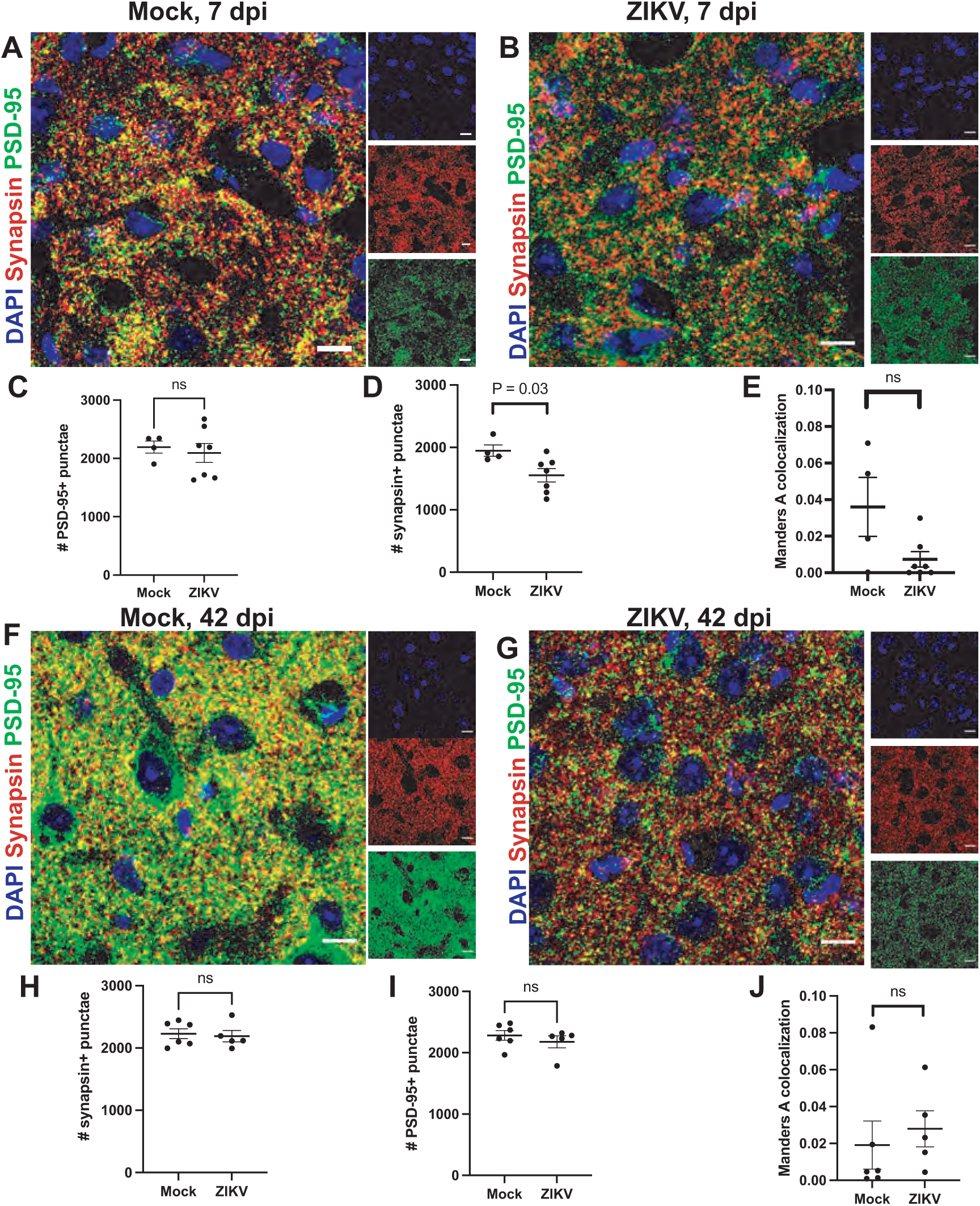
FC in ZIKV infection is associated with decreased of pre-synaptic markers during acute infection and recovered synapses at 42 days post infection. Representative synapse staining in the somatosensory cortex of (A) mock and (B) ZIKV-infected mice at 7dpi. (C) Synapsin, a pre-synaptic marker, is decreased in the somatosensory cortex at 7 dpi. (D) PSD-95, a post-synaptic marker, is stable in mock and infected mice. (E) Colocalization of synapsin and PSD-95 is not significantly different. Images taken at 63x; scale bars: 10 μm. (D) Quantification of synapse markers showing mock (n = 4) and Zika-infected (n= 7) mice. Data pooled from 2 independent experiments. Representative synapse staining in the somatosensory cortex of (F) mock and (G) ZIKV-infected mice at 7dpi. (H) Synapsin, a pre-synaptic marker, and PSD-95 (I), a post-synaptic marker do not differ between mock and infected mice after recovery. Images taken at 63x; scale bars: 50 μm. (J) colocalization between pre- and post-synaptic markers is not difference. Quantification of synapse markers showing mock (n = 6) and Zika-infected (n= 5) mice. Data pooled from 2 independent experiments.

## Discussion

Although the importance of immune cell infiltration and activation during acute viral infection is well-established, its effect on CNS recovery after viral clearance is poorly understood. In this study, we demonstrate that loss of cortical FC is associated with neuronal injury and synapse loss during acute ZIKV encephalitis. Via repeated FC assessment, we also determined that recovery of cortical FC is positively correlated with myeloid cell activation during the acute infection, suggesting that persistent inflammation may play a role in brain recovery from viral infection. Use of MA-ZIKV, which causes severe injury of the hippocampal formation, also reveal discoupling between cortical FC recovery and hippocampal-based learning.

Recent FC research has uncovered the complexity of the acquired brain network data by demonstrating that functional networks reflect not only neuron health, but also the health of synapses, glia and microglia that support brain networks^34^. In this study, we evaluated resting state cortical FC in conjunction with immunohistochemical analyses to determine if functional neuroimaging might correlate with neurocognitive findings as a noninvasive assessment of cognitive status during acute infection with, and recovery from, ZIKV encephalitis. In prior studies, ZIKV-Dak strain has been shown to primarily target the hippocampus. Here, we selected a highly virulent, mouse adapted strain of Zika-Dak (ZIKV-Dak-MA) for our study, which exhibits widespread infection throughout the brain, including the cerebral cortex, with highest viral titers at 7 dpi and complete viral clearance by 15 dpi. This strain was also observed to induce severe destruction of the hippocampal formation. Consistent with this, we found that, compared to mice infected with ZIKV-Dak, mice infected with ZIKV-Dak-MA were completely unable to learn a spatial memory task assessed via the Barnes maze. We wondered if widefield optical imaging could detect this persistent cognitive deficit as alterations in cortical FC. While we found that cortical FC decreased in the somatosensory cortex during acute infection (7 dpi), it appeared to largely recover at 42 dpi, despite hippocampal obliteration.

We also performed immunohistochemical studies in order to correlate FC findings with significant alterations in inflammatory markers during acute ZIKV encephalitis. Our analyses revealed not only decreased neurons and increased GFAP- and Iba1- positive cells in the somatosensory cortex, but similar trends in the retrosplenial cortex, which did not exhibit FC deficits. These data indicate these acute responses to infection correlated with FC findings. Similarly, we found that neuronal numbers appeared to recover, but numbers of GFAP+ cells remained increased in the somatosensory cortex at 42 dpi. Similar to the inflammation signaled by increased GFAP+ cells at 42 dpi in the cortex, we detected persistent myeloid cell activation with increased Iba1- positive cells throughout the hippocampus at 42 dpi, and severe injury, with ablation of neurons in the CA3 region. These data support our findings that i.c. ZIKV-Dak-MA-infected mice are unable to learn the Barnes maze after viral clearance and recovery. Nevertheless, these findings demonstrate that functional networks are driven by synaptic function during the course of ZIKV infection and recovery as measured by widefield optical imaging. This study also contributes to and supports the growing body of literature to suggest contributions of neurons, glia, microglia and synapses to resting state-FC networks^34, 36–38^.

In addition to the primary findings of the study, we also found high average delta power in the Zika-infected mice at 7 dpi, which returned to levels close to that of mock-infected mice at 42 dpi. Previous studies have shown that increased Delta power is correlated with poor outcome in patients with stroke, for example^39^, and this is directly related to neuronal metabolism. This indicates that delta power is also inversely related to the health of brain networks and provides different information about cognition from that found in interhemispheric connectivity measures. This would be consistent with our findings that mice infected with ZIKV do, in fact, have long term cognitive sequelae, even though their interhemispheric cortical connectivity recovers.

The findings in this study may have some limitations. We found that when we repeatedly imaged the same mice at 7 dpi and 42 dpi, not only did ZIKV-infected mice exhibit similar FC compared to mock-infected mice at 42 dpi, but mock-infected mice demonstrated a decrease in FC from 7 dpi to 42 dpi. As mice were infected and imaged as young adults (8-9 weeks old), and imaged again at 42 dpi (15 weeks old), it is possible that aging contributed to our findings at the recovery time-point. Previous studies have shown that connectivity does change over the lifespan^40, 41^, but there is little functional neuroimaging data evaluating the ages used in our study. This may be an area for future investigation.

While excellent two-dimensional spatial resolution and time-series data is captured with widefield optical imaging, another known limitation of widefield optical imaging is the depth of resolution due to the effects of light scattering and absorption. Previous studies have shown persistent alterations in cortical FC in mouse models of glioma growth^27^, stroke^28^, and Alzheimer’s Disease (AD)^29^, which were presumed to include hippocampal-cortical networks. Our model indicates, however, that alterations in cortical FC do not necessarily reflect hippocampal-cortical networks, as recovery of cortical FC occurs despite severe hippocampal injury shown by both immunohistochemical and behavioral approaches. Notably, we did not observe FC alterations in the retrosplenial cortex, which includes both cortical and hippocampal networks.

In summary, our data demonstrate that cortical FC decreases during acute ZIKV encephalitis due to loss of presynaptic termini, which improves after recovery from infection. This connectivity and synapse recovery correlated with the degree of myeloid cell activation, implicating neuroinflammation in the recovery of cognitive networks in the brain after ZIKV infection. This data has important implications for immunomodulatory therapies in post-infectious neurocognitive recovery, as it suggests that not all neuroinflammation is detrimental. Thus, selective immunomodulatory therapies that maintain myeloid cell activation may actually promote recovery. Furthermore, although we saw similar trend in astrocytes that did not reach significance, activated astrocytes may contribute to these effects. Future studies are needed to dissect the individual contributions of myeloid cells and astrocytes in cortical synaptic recovery.

## Materials and Methods

### Animals

A total of 51 two month old mice (mock, N=25, ZIKV, N=26) consisting of 24 female and 21 male mice total (N=12 female, N=10 male mock and N=12 female, N=11 male ZIKV mice) *Thy1*-GCaMP6f mice were used in the present study (Jackson Laboratories Strain: C57BL/6J-*Tg(Thy1-GCaMP6f)GP5*.*5Dkim*; stock: 024276). These mice express the protein GCaMP6f in excitatory neurons, primarily in cortical layers ii, iii, v, and vi^26^. All studies were approved by the Washington University School of Medicine Animals Studies Committee and follow the guidelines of the National Institutes of Health’s Guide for the Care and Use of Laboratory Animals.

To evaluate for effects of intracranial needle insertion, mock mice are compared to mice injected peripherally with PBS 72 hours prior (N=8, all female, 3-7 months old) (Suppl. Fig 3). These mice underwent typical surgical preparations as described below, with additional bilateral EEG implantation (not used in the present study).

### ZIKV infection

The ZIKV mouse adapted (MA) Dakar strain utilized for intracranial infections was obtained from M. Diamond at Washington University in St Louis. The MA Dakar strain of ZIKV was obtained by passage of the original Dakar strain of ZIKV through a *Rag* -/- mice^31^, resulting in a strain of ZIKV that was found to replicate more efficiently in the mouse brain than the parent strain. Mice were deeply anesthetized and intracranially administered 1 × 10^4^ plaque-forming units (p.f.u.) of ZIKV MA Dakar. Viruses were diluted in 10 µl of 0.5% fetal bovine serum in Hank’s balanced salt solution (HBSS) and injected into the midline of the brain with a guided 29-guage needle. Mock-infected mice were intracranially injected with 10 µl of 0.5% fetal bovine serum in HBSS into the midline of the brain with a guided 29-guage needle.

### Surgical windowing

Prior to imaging, a plexiglass optically transparent window was implanted with translucent dental cement (C&B-Metabond, Parkell Inc., Edgewood, New York) following a midline incision and clearing of skin and periosteal membranes. The window covered the majority of the dorsal cortical surface and provided an anchor for head fixation and allowed for chronic, repeatable imaging^42^.

### Immunohistochemistry

After imaging at either 7 dpi or 42 dpi on the same day, Mice were anesthetized and perfused with ice-cold Dulbecco’s PBS (Gibco) followed by ice-cold 4% paraformaldehyde (PFA). Brains were post-fixed overnight in 4% PFA followed by cyroprotection in 30% sucrose (three exchanges for 24 h), then frozen in OCT compound (Fischer). Coronal tissue sections (10 µm) were washed with PBS and permeabilized with 0.1–0.3% Triton X-100 (Sigma-Aldrich). Nonspecific antibodies were blocked with 5% normal goat serum (Sigma-Aldrich) at room temperature for 1 h. To reduce endogenous mouse antibody staining when detecting ZIKV, a Mouse on Mouse kit (MOM basic kit, Vector) was used as per the manufacturer’s instructions. Slides were then incubated in primary antibody (described below) or isotype-matched IgG overnight at 4 °C. Following washes in PBS, slides were then incubated in secondary antibodies at room temperature for 1 h, and nuclei were counterstained with 4,6-diamidino-2-phenylindole (DAPI; Invitrogen). Coverslips were applied with ProLong Gold Antifade Mountant (Thermo Fisher). Immunofluorescent images were acquired using a Zeiss LSM 880 confocal laser scanning microscope and processed using software from Zeiss. Immunofluorescent signals were quantified using the software ImageJ and using in-house written scripts using Matlab (Natick, MA). Antibodies used can be found in supplemental material.

### RNAscope in situ hybridization

Tissue was prepared similarly to that used for immunohistochemistry. RNAscope 2.5 HD Assay-Brown was performed as per the manufacturer’s instructions. Probes against and ZIKA mRNA (ACD) were used. Viral tropism was analyzed using a ZEISS Axio Imager Z2 fluorescence microscope.

### Fluorescence and Optical Intrinsic Signal (WOI) Imaging

Mice were head-fixed in a stereotaxic frame and body secured in a black felt pouch for imaging. Sequentially firing LEDs (Mightex Systems, Pleasanton California) passed through a series of dichroic lenses (Semrock, Rochester New York) into a liquid light guide (Mightex Systems, Pleasanton California) that terminated in a 75mm f/1.8 lens (Navitar, Rochester New York) to focus the light onto the dorsal cortical surface. LEDs consisted of 470nm (GCaMP6f excitation), 530nm, 590nm, and 625nm light. An sCMOS camera (Zyla 5.5, Andor Technologies, Belfast, Northern Ireland, United Kingdom) coupled to an 85mm f/1.4 camera lens (Rokinon, New York New York) was used to capture fluorescence/reflectance produced at 16.8 Hz per wavelength of LED. A 515nm longpass filter (Semrock, Rochester New York) was used to discard GCaMP6f excitation light. Cross polarization (Adorama, New York New York) between the illumination lens and collection lens discarded artifacts due to specular reflection. The field-of-view (FOV) recorded covered the majority of the convexity of the cerebral cortex (∼1.1cm^2^), extending from the olfactory bulb to the superior colliculus. All imaging data were acquired as 5-min runs and binned in 156x156 pixel^2^ images at approximately 100 µm per pixel. All 45 mice were imaged at 7dpi while a smaller cohort (mock, N=11, ZIKV, N=8) were imaged post recovery at 42dpi.

### Behavioral testing

Animals underwent open-field and Barnes maze testing as previously described13. Briefly, mice were given 3 min to explore the Barnes maze platform and to find the target hole. If a mouse did not find the target hole within the test period, it was gently guided in the hole. At the end of a trial, the mice remained in the target hole for 1 min before being placed back into their home cage between trials. Each mouse received two trials per day over the course of five consecutive days. The number of errors (or number of nose pokes over non-target holes) was recorded. Before Barnes maze testing, animals performed the OFT as a measure of exploratory behavior. Each mouse was given 5 min to explore an empty arena before being placed back into its home cage. Both the Barnes maze platform and the open-field arena were decontaminated with 70% ethanol between trials and/or mice. All trials were video recorded using a camera (Canon Powershot SD1100IS) and scored by experimenters blinded to the treatment conditions.

### Statistical analysis for histochemical assays

Unpaired Student’s t-test or one-way or two-way analysis of variance (ANOVA) with appropriate post-test to correct for multiple comparisons, in which the means of all groups were compared pairwise, were performed as indicated in the figure legends. Prism 7.0 (GraphPad Software) was used for generating graphs and statistical analyses. P values of ≤0.05 were considered significant. To determine sample sizes for virological and immunological studies, a power analysis was performed using the following values: probability of type I error = 0.05, power = 80%, fivefold hypothetical difference in mean, and population variance of 25-fold (virological studies) or 12-fold (immunological studies). Sample sizes for behavioral testing experiments were predetermined by the power analysis. For all experiments, animals were randomly assigned to mock or MA- ZIKV-Dak infection. Investigators were blinded to group allocation during data collection and analyses.

### Imaging data processing

Image processing followed methods previously described^25, 43^ and briefly summarized here. Images were spatially downsampled to 78x78 pixel^2^ and a frame of ambient baseline light levels was subtracted from the time series data. Data were temporally downsampled by a factor of 2 and then spatially and temporally detrended. Data were affine-transformed to common Paxinos atlas space and pixel-wise time traces were mean normalized. Frames corresponding to reflection data produced by the 530nm, 590nm, and 625nm light were used to solve the modified Beer Lambert law to yield fluctuations in oxygenated and deoxygenated hemoglobin. Frames corresponding to fluorescence data were corrected by approximating hemoglobin absorption of the excitation and emission light. All data were spatially smoothed with a 5x5 Gaussian filter. The global signal was regressed from the time series data, and data were filtered with a 0.4-4.0Hz Butterworth bandpass filter.

### Image data analysis

For functional connectivity (FC) analysis we calculated the Pearson correlation coefficient between the time traces from two different regions:

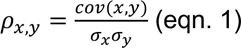

Where 𝜌_𝑥,𝑦_ is the Pearson correlation coefficient between time traces 𝑥 and 𝑦, 𝑐𝑜𝑣(𝑥, 𝑦) is the covariance between time traces 𝑥 and 𝑦, and 𝜎_𝑥_, 𝜎_𝑦_ are the standard deviations within time trace 𝑥 and time trace 𝑦, respectively. Bilateral FC maps represent Pearson correlation coefficients calculated between pixel-wise time traces on the left hemisphere and their corresponding symmetrical right hemisphere pixel-wise time trace. Power spectral analysis of the GCaMP6f data was performed on 10 second segments by applying a Hann window and an FFT (squared to obtain power).

### Statistics

A challenge with analyzing the statistical significance in functional imaging is managing the multiple comparisons problem. Here, we used a cluster size-based method that leverages the spatial connection between pixels, and credits large clusters as having more statistical significance than small clusters with the same peak *t*-value. More specifically, we used a cluster size-based thresholding method to analyze FC maps, and to ensure the family-wise error rate of any thresholded map did not exceed *p*=0.01^44^.

## Acknowledgments

This work was supported by the National Institutes of Health grant numbers R01NS104471, R35 NS122310 (to RSK), R01NS099429 and R01NS090874 (to JPC), F31NS110222 (to RMR), (to SCA), and F30AG061932 (to LMB).

