## Supplemental Data for "Myeloid cell activation during Zika virus encephalitis predicts recovery of functional cortical connectivity"

### Supplementary Data

**Supplemental Figure 1.** Evaluation of mouse adapted (MA) strain of Zika virus. (A) Survival curves and (B,C) weights and encephalitis scores comparing MA and parent Dakar strain. (D-G) Viral titer levels at 3, 7, and 15 days after intracranial infection. In situ hybridization on sagittal mouse brain sections of (H) parent and (I) mouse-adapted Zika strains. Behavioral testing data showing (J) Barnes maze and (K) open field and data comparing mock, parent strain Dakar and MA Dakar infected mice at 42 days post infection

**Supplemental Figure 2. ZIKV infection is associated with cellular changes in the hippocamapus during acute infection and severe hippocampal injury after recovery from infection.** Representative CA3 regions from mock (A) and ZIKV-infected (B) mice at 7 dpi. There is a trend toward decreased NeuN+ neurons (C) and significantly increased GFAP+ (D) and Iba1+ (E) cells in the CA3 at 7 dpi. At 42 dpi, representative images of the mock-infected CA3 (F) compared to ZIKV-infected (G) CA3 shows severe injury to the ZIKV-infected CA3. and decreased pre-synaptic marker, synapsin, in the somatosensory cortex. (A) Representative figures demonstrating decreased numbers of neurons with associated increased monocytes and astrocytes in the cornu ammonis 3 (CA3) region of the hippocampus in Zika-infected (bottom) mice compared to mock-infected (top) mice. Images taken at 20x.

**Supplemental Figure 3. FC deficit align most strongly with the somatosensory cortex.** Cortical Paxinos atlas parcellation adapted by White et al.[20] with cortical regions defined and deficit from bilateral FC outlined in white.

**Supplemental Figure 4. ZIKV infection effects hemodynamics more than calcium, in a sex-independent manner.** Histogram of Pearson correlation values in the average bilateral FC maps from Figure 1 showing the whole FOV (top) and only the regions in the cluster-defined ROI (bottom) for A) delta calcium and B) infraslow hemoglobin. Average (mock N=22, ZIKV N=23) bilateral FC over the FOV separated by sex at 7dpi for C) delta calcium and D) infraslow hemoglobin. Average (mock N=11, ZIKV N=8) bilateral FC over the FOV separated by sex at 42dpi for C) delta calcium and D) infraslow hemoglobin. Significance testing by a two-sample t-test.

**Supplemental Figure 5: Mock infected FC is relatively unaffected by cranial needle injection.** Average pixel-wise (PBS N=8, mock N=22) bilateral correlation maps (top row) and standard deviations (middle row) across mice. Pixel-wise two-sample *t*-test (bottom row, left) and thresholded image (bottom row, right) for $p<0.01$ by a cluster-based thresholding method.

**Supplemental Figure 6.** ZIKV infection weakens multiple network connections using hemoglobin infraslow dynamics. A,B) Average (mock N=22, ZIKV N=23) FC matrices displaying the Pearson correlation between network seeds shown on the x and y-axis, as well as the C) difference in Pearson correlation between the mock and ZIKV matrices. D) Network-wise two-sample *t*-test, and E) corresponding result highlighting regions with a $p$-value<0.05 (uncorrected for multiple comparisons). F) Matrices are thresholded to display $p$-values below the Bonferroni threshold for significance (two-sample *t*-test).

**Supplemental Figure 7: A smaller ZIKV sample size yields the same somatosensory based FC deficit at 7dpi in delta calcium.** Top row) Average (mock, N=11, ZIKV, N=8) pixel-wise bilateral FC maps across mice. Bottom row) Pixel-wise two-sample *t*-test (left) and thresholded image (right) for $p<0.01$ by a cluster-based thresholding method.

**Supplemental Figure 8: A smaller ZIKV sample size yields the same global FC deficit at 7dpi in infraslow hemoglobin.** Top row) Average (mock, N=11, ZIKV, N=8) pixel-wise bilateral FC maps across mice. Bottom row) Pixel-wise two-sample *t*-test (left) and thresholded image (right) for $p<0.01$ by a cluster-based thresholding method.

**Supplemental Figure 9. Acute ZIKV increases global delta power.** Histogram displaying pixel-wise GCaMP6 delta power across the whole FOV at 7 dpi (left) and 42 dpi (right). Delta power increased during acute infection and recovers to healthy delta power after recovery from ZIKV infection.

**Supplemental Figure 10. FC at 7 dpi does not predict FC at 42 dpi, but degree of myeloid cell activation does.** Scatter plots of FC at 7 dpi vs. FC at 42 dpi (A) and Iba1-positive quantification vs. 42 dpi FC (B), demonstrating that FC at 7 dpi and 42 dpi do not correlate, but number of Iba1 positive cells in ZIKV-infected mice does correlate with FC at 42 dpi.

**Supplemental Figure 11. FC decreases with age in healthy mice**. FC for all mock and ZIKV-infected mice at 7 dpi and 42 dpi, illustrating that even in mock mice, FC decreases between 7 dpi and 42 dpi.

### Supplemental Figure 1

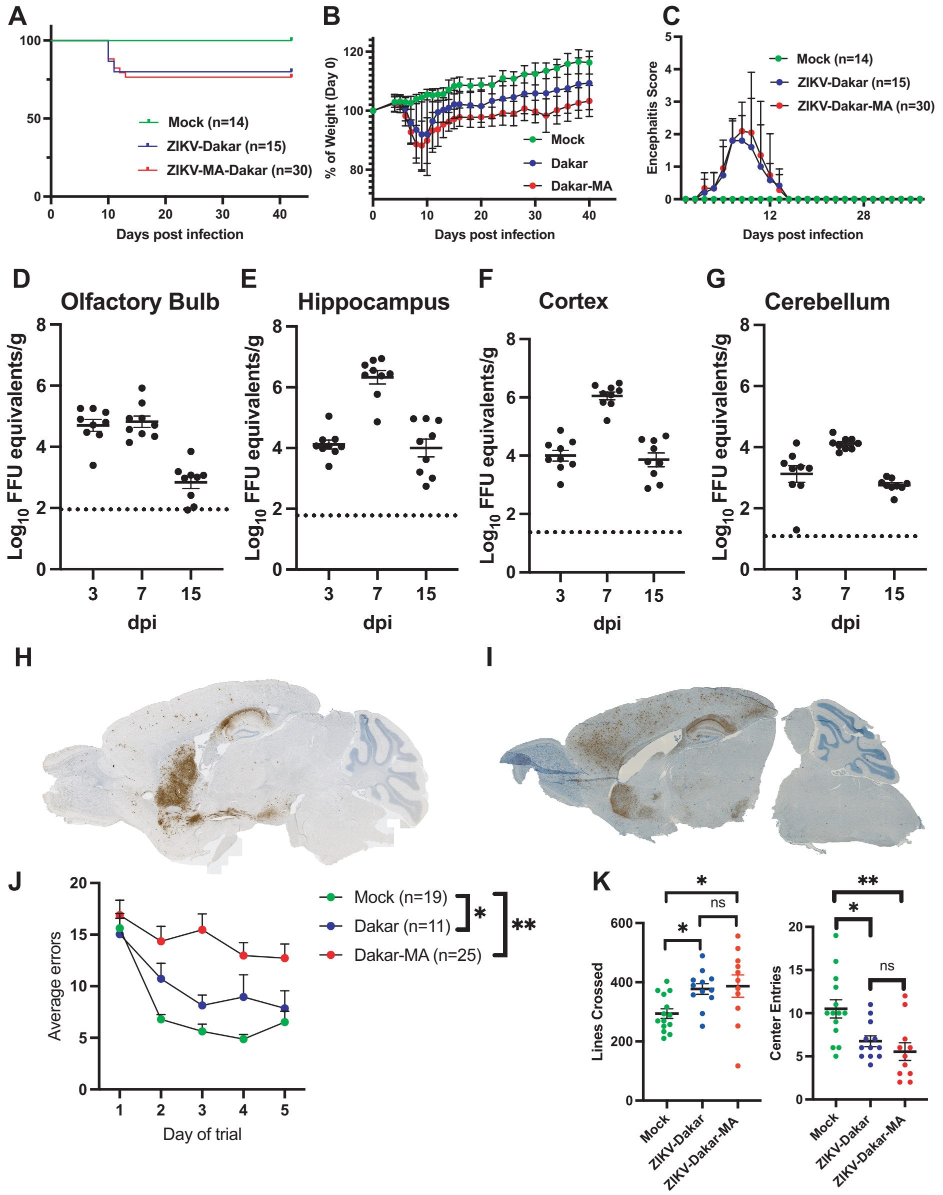

### Supplemental Figure 2

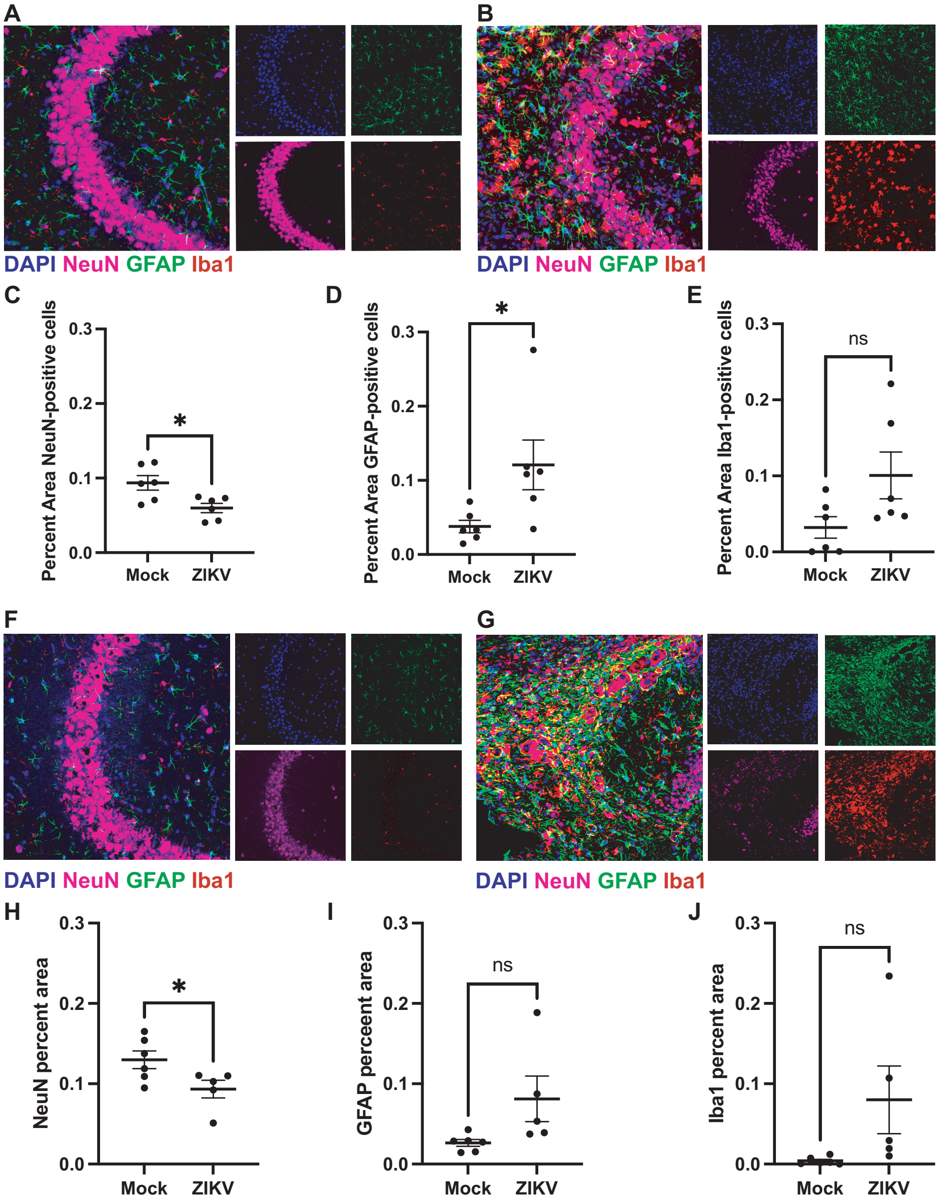

**Supplemental Figure 3**

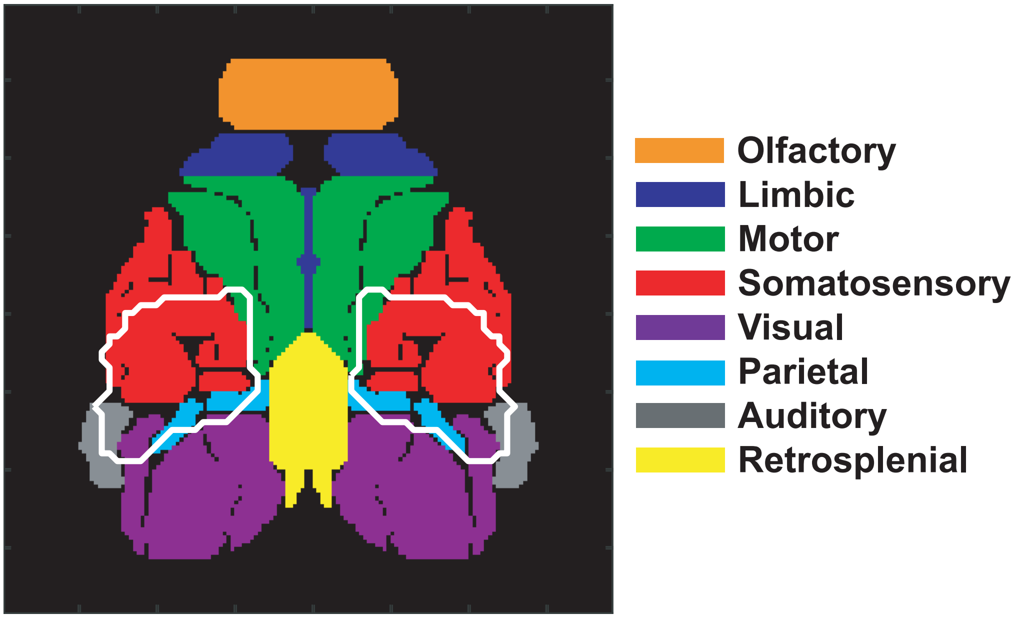

### Supplemental Figure 4

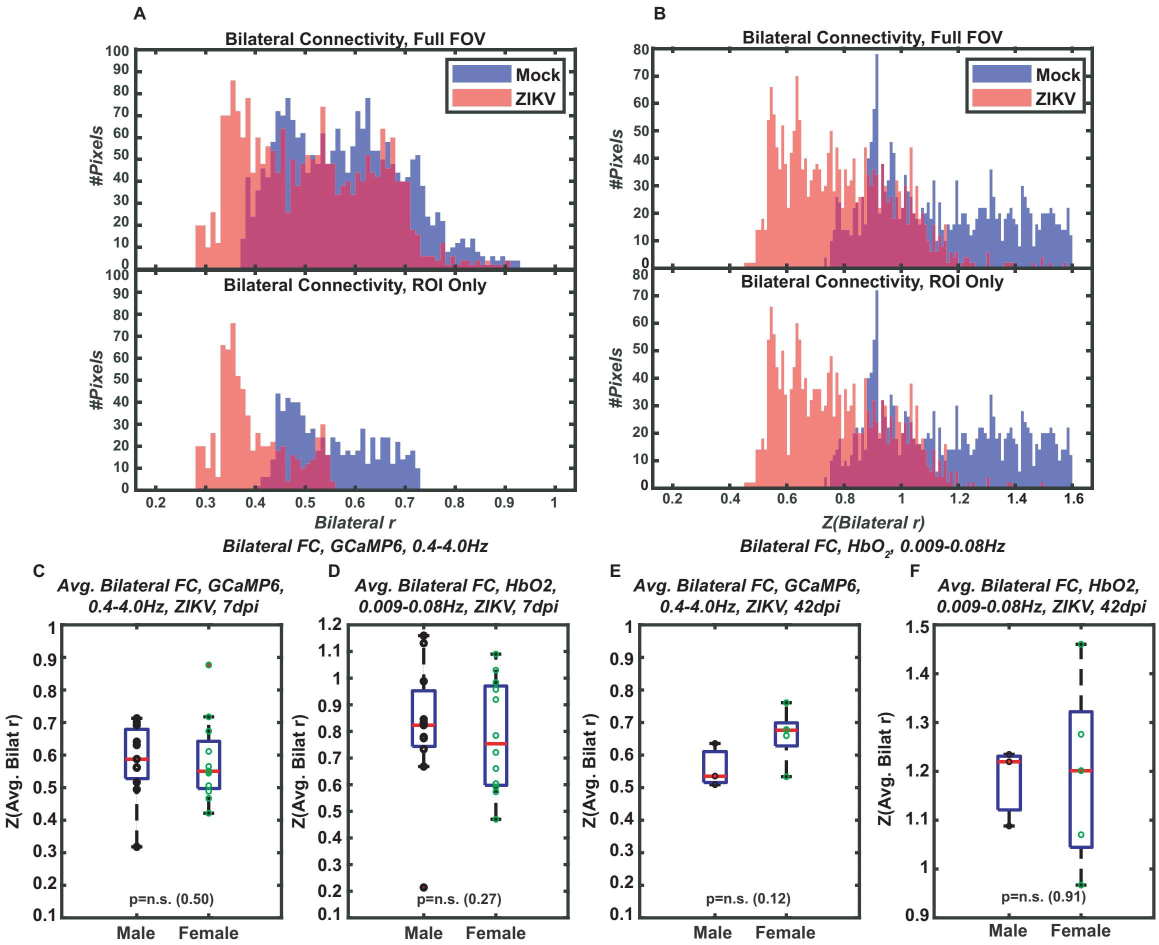


Supplemental Figure 4

### Supplemental Figure 5

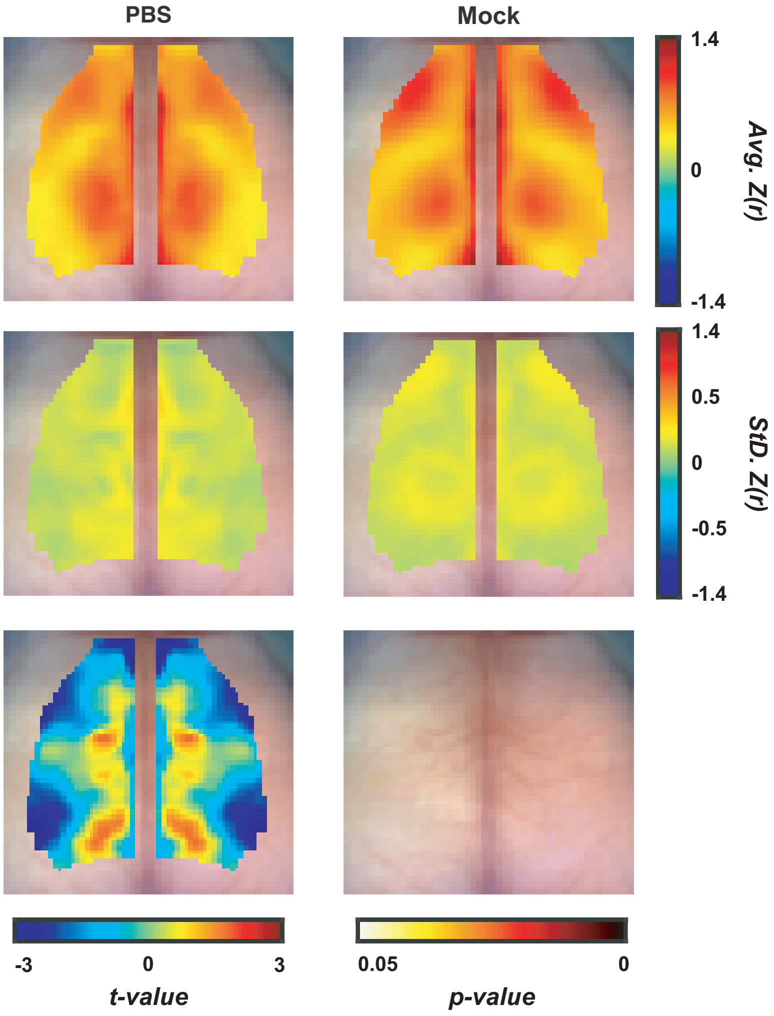


*Bilateral FC, GCaMP6, 0.4-4.0Hz*

### Supplemental Figure 6

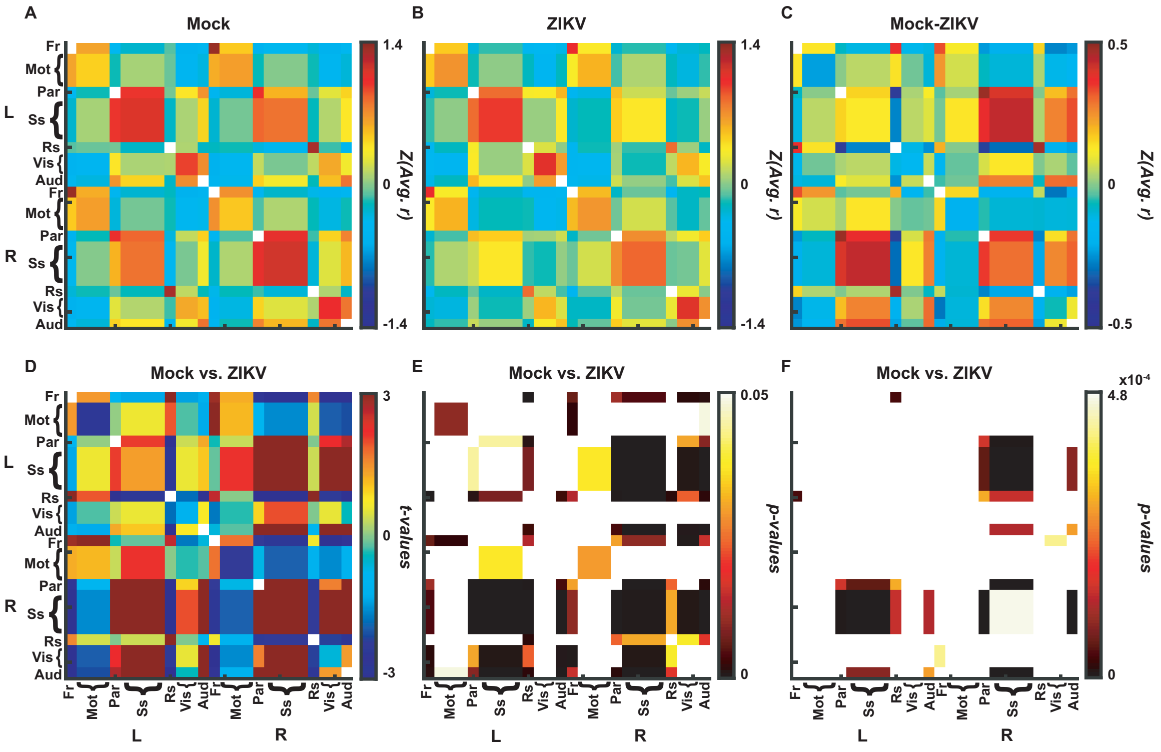


*FC, $HbO_2$, 0.009-0.08Hz*

### Supplemental Figure 7

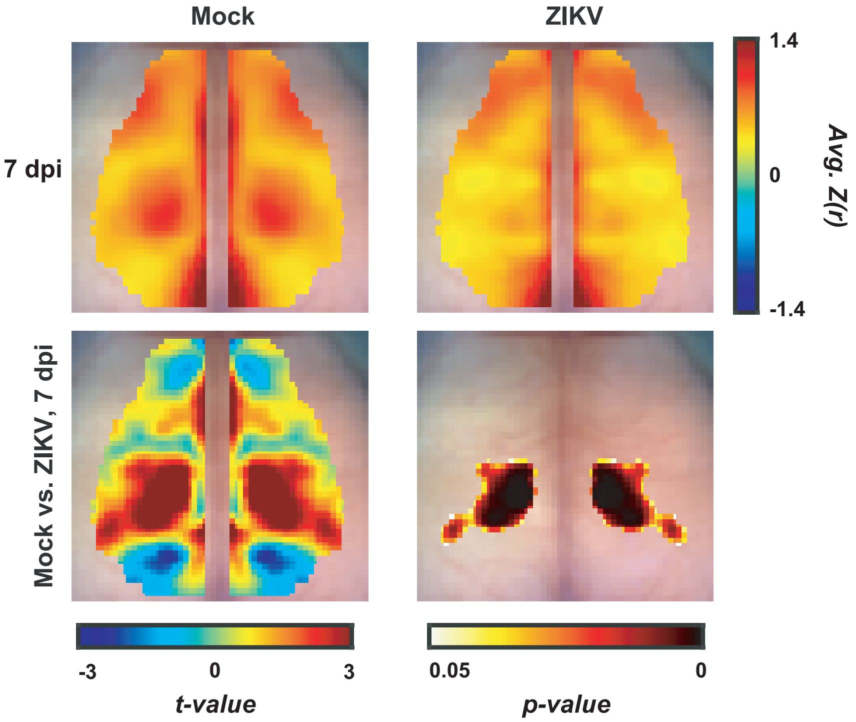


*Bilateral FC, GCaMP6, 0.4-4.0Hz*

**Supplemental Figure 8**

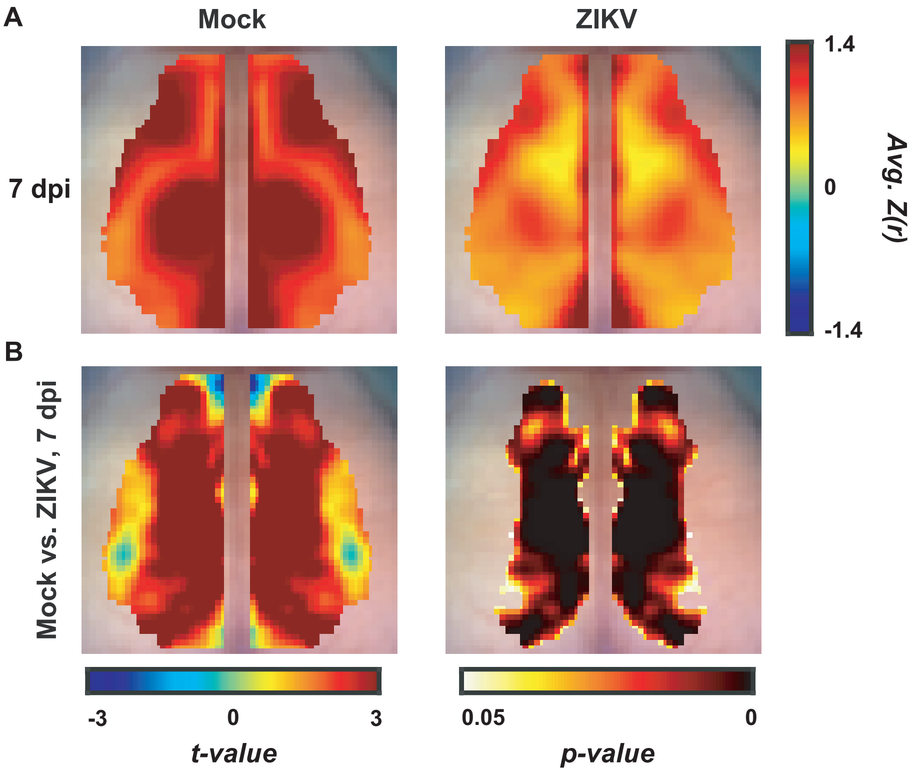


*Bilateral FC, $HbO_2$, 0.009-0.08Hz*

### Supplemental Figure 9

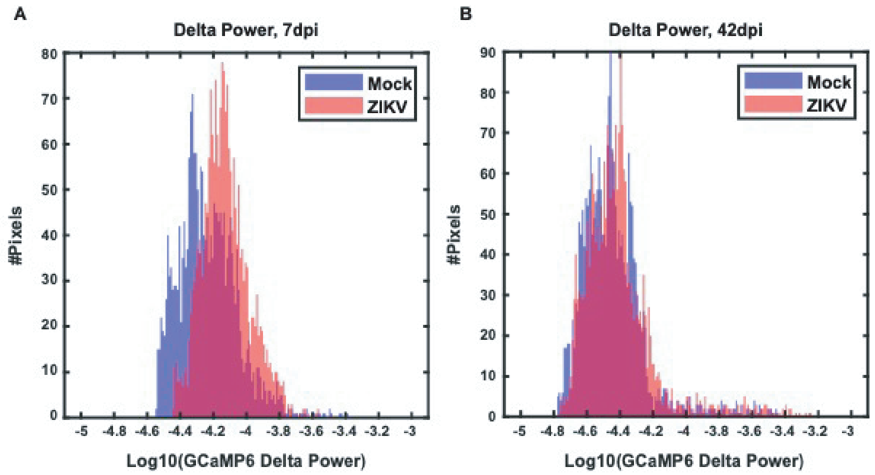


### Supplemental Figure 10

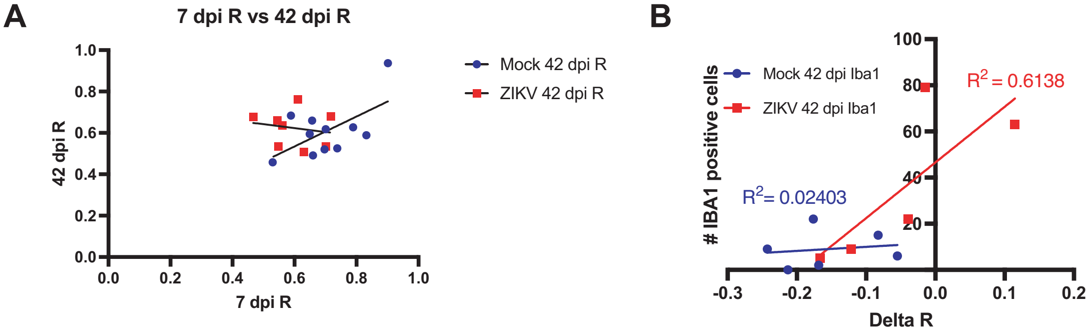


### Supplemental Figure 11

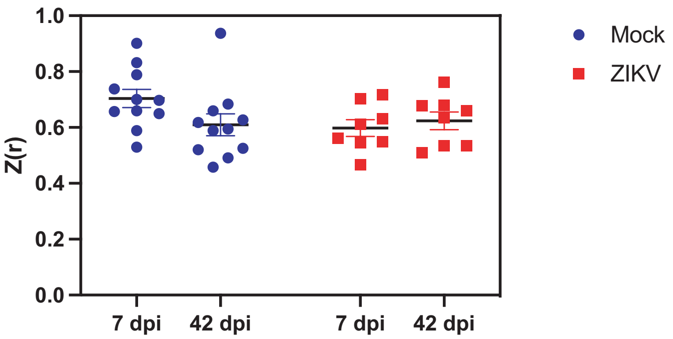
